# Absolute pitch can be learned by some adults

**DOI:** 10.1101/325050

**Authors:** Stephen C. Van Hedger, Shannon L.M Heald, Howard C. Nusbaum

## Abstract

Absolute pitch (AP), the rare ability to name any musical note without the aid of a reference note, is thought to develop in an early critical period of development. Although recent research has shown that adults can improve AP abilities in a single training session, the best learners still did not achieve note classification performance comparable to performance of a genuine AP possessor. Here, we demonstrate that genuine AP levels of performance can be trained in eight weeks for some adults, with the best learner passing all measures of AP ability after training and retaining this knowledge for at least four months after training. Alternative explanations of these positive results, such as improving accuracy through adopting a slower, relative pitch strategy, are not supported based on joint analyses of response time and accuracy. The post-training AP assessments were extensive, totaling 204 notes taken from eight different timbres and spanning over seven octaves. These results clearly demonstrate that explicit perceptual training in some adults can lead to AP performance that is behaviorally indistinguishable from AP that results from childhood development. Implications for theories of AP acquisition are discussed.

## 1. Introduction

Absolute pitch (AP) is the rare ability to name any musical note without the aid of a reference note ^1–3^. The question of how individuals acquire this ability continues to be a matter of debate. The most widely accepted theory is that “genuine” AP ability can only manifest within an early period of development (the *critical period* theory) ^4,5^. In part, the *critical period* theory of AP is bolstered by the lack of conclusive evidence that AP can be learned by post-critical period adults ^6–9^ as well as a more recent study suggesting that there is a need to re-open the critical period for learning AP with a pharmacological intervention ^10^.

Here, we directly test the hypothesis of a critical period for AP acquisition through intensive AP training in a post-critical period adult sample. One key difference between this study and previous AP training studies is that participants were selected specifically based on auditory working memory (WM) abilities. Recent research shows that individual differences in auditory WM predict how well adults can learn AP categories from a single training session, although performance was well below thresholds typically used to designate genuine AP ability ^11^. Yet, given this observed relationship between auditory WM and AP learning, we asked whether providing substantially more AP training for adults with high auditory WM abilities can produce performance levels comparable to the genuine AP listener, specifically regarding retention of learning (over a timescale of months) and accurate performance across a range of instrumental timbres and octaves. This type of demonstration, even in a single adult without prior AP ability, would inform our understanding of the *critical period* theory of AP.

As an alternative to the *critical period* theory, AP can be conceptualized as an auditory skill ^12^ that is shaped by both short- and long-term experiences ^17,18^. This view (which we refer to as the *skill acquisition* theory) supports the prediction that listeners should be able to improve AP performance through explicit perceptual training at any age, with at least some individuals exceeding typical thresholds for genuine AP inclusion post-training. If, however, individuals are only able to modestly improve in AP categorization, with no individual exceeding AP thresholds post-training, this suggests that there might be fundamental limits that constrain performance as would be expected under a critical period framework. Thus, the question of whether a post-critical period adult can learn AP has important implications for understanding individual differences in AP performance, the underlying mechanisms of AP, and the importance of environmental factors on developing and maintaining AP ability.

Six adult participants completed an eight-week AP training regime. Prior to training, participants completed tests of auditory WM, short-term memory (STM), and AP ability. The eight-week AP training program was divided into two phases lasting four weeks each. In both phases of training, participants had to complete three training protocols four times each week, which amounted to approximately 4 hours of training per week (32 hours in total). Every week, participants also completed an AP test in which isolated notes were categorized without feedback. In the second phase of training, we added an additional test that required participants to label the key signature of a presented melody without feedback. At the end of the eight-week training program, participants were retested on the same AP tests that were administered prior to training, in addition to an AP test not previously administered but widely used in prior research. Approximately four months later, we retested participants on the same AP tests as those administered post-training to assess whether learning remained stable.

## 2. Method

### 2.1 Participants

Six participants (three female, three male) participated in the experiment (*M* = 23.33, *SD* = 2.94 years old, age range: 18-26). None of the participants self-reported possessing AP. All participants provided written consent and were compensated for their participation in the experiment.

### 2.2 Task Description and Materials

#### 2.2.1 AP Training Program

The absolute pitch training program was eight weeks in duration. There were two phases to the experiment (each lasting four weeks). Both phases consisted of three training programs that participants had to complete four days every week. Participants were tested every Friday, meaning that participants could complete their weekly training programs Saturday through Thursday.

##### First Phase

Two of the First Phase training programs were meant to emphasize speed in classifying absolute pitches, while the third was meant to emphasize accuracy. In the first program, nicknamed “Simple Speed” (SS), participants would see a note name presented in the center of the screen for 1500ms (e.g., C). This was the target note for the trial. During the presentation of the note name, participants heard a sung version of the target note, which was enunciated with the category label. After participants heard the sung target note, they heard a string of 16 notes, with an inter-note-interval of 2250ms. Within these 16 notes, 25% (4 of 16) were the target note, while 75% (12 of 16) were non-target notes. Target notes were randomly interspersed with non-target notes. Participants had to press the spacebar as quickly as possible whenever they heard the target note. Participants had 1750ms to respond to target notes. All notes were taken from a C-major scale (i.e., white keys only) spanning a one-octave range (C [4] to B [4]). Non-target notes were also taken from a one-octave range and consisted of only white keys. The notes were synthesized with a piano timbre. Participants received feedback after each note. There was a 1500ms rest period between trials. Each of the seven target notes (C, D, E, F, G, A, and B) were presented twice, for a total of 14 trials during each session.

The second program, nicknamed “Complex Speed” (CS), followed the same general procedure as SS, with the following differences. First, the presentation of the 16 notes was faster (1750ms), meaning that participants had to respond more quickly (within 1250ms). Second, the notes (both target and distractor) could come a piano, flute, or guitar timbre. Third, the octave range was expanded relative to the SS task (C [3] to B [5]).

The third program, nicknamed “Accuracy Training” (AT), emphasized correct note category identification over speed of identification. There were two blocks, consisting of 48 trials each (96 trials total). The general procedure was the same for both blocks. On each trial, participants would hear an isolated note, which was presented simultaneously with a response screen. The response screen displayed all twelve note-category options, arranged in a pitch wheel. Participants then had an unlimited amount of time to click on the corresponding note name with the mouse. After each response, participants would receive feedback, in which the correct note name was highlighted and participants reheard the note. The first block consisted of piano notes, spanning a two-octave range (C [4] to B [5]). Unlike the SS and CS tasks, all 12 categories were explicitly trained. Since there were 24 total piano notes (12 note categories × 2 octaves), each note was presented twice in a random order. The 48 notes in the second block were randomly selected from 96 possible notes (24 piano, 24 cello, 24 clarinet, and 24 harpsichord). Participants heard 1000ms of noise between trials to minimize the influence of relative pitch information in selecting the correct absolute pitch.

Every Friday, participants would have to complete a weekly test (WT) of AP ability. The WT was similar to the AT task, with two exceptions. First, participants did not receive feedback. Second, the second block was expanded to 120 possible notes (24 piano, 24 cello, 24 clarinet, 24 harpsichord, and 24 square wave).

##### Second Phase

We replaced the SS and CS tasks after Week 4 with a more difficult speeded task, nicknamed “Hypercomplex Speed” (HS). The HS task followed the same general procedure as the SS and CS, with the following exceptions. First, on each trial, participants heard a string of 32 notes, with an inter-note-interval of 1500ms. Within these 32 notes, 12.5% (4 of 32) were the target note, while 87.5% (28 of 32) were non-target notes. Second, the response decision window was reduced to 1000ms. Third, both the target and the non-target notes were taken from all 12 note categories (sampling from all 160 note stimuli described in Table S1).

We also introduced a novel task during the Second Phase nicknamed “Name That Key” (NTK). The NTK task followed the same general procedure as the AT task. Each session, participants heard 15 music recording excerpts, which were randomly selected from a database of 300 total recordings. Similar to the AT task, participants had to decide on the key signature of the recording by clicking on a pitch wheel with the 12 pitch categories arranged around a circle. After each selection, participants would see a specific feedback screen, in which the correct note name was highlighted on the pitch wheel. Additionally, participants heard the tonic note of the key signature (e.g., C) played during feedback. Participants heard 1000ms of noise between trials. Participants also completed the AT task during the Second Phase, which was identical to the version presented in the First Phase.

Every Friday, participants would have to complete the same WT from the First Phase, in addition to a new test, nicknamed the “Name That Key Test” (NTKT). The NTKT required participants to judge the key signature of 12 folk songs. During the task, each pitch class was represented by a key signature one time (i.e., one folk song played in C, one folk song played in C#, etc.), and feedback was not provided. The randomized assignment of folk melody to key signature was hard coded into the script, as we did not want any given folk song to play in the same key signature across weeks.

The isolated musical note stimuli used in training were created using Reason Music Production Software (Propellerhead: Stockholm, Sweden). Musical notes were sampled from real instruments and were digitized at a 44.1 kHz sampling rate with 16-bit depth, were 1000ms in duration, and were root mean square normalized to a level of −5 dB FS. We used a total of 160 notes (from seven instrumental timbres) throughout training. Details of the specific notes used can be found in Table S1. For the music recording stimuli used in the NKT task, we recorded 15-second excerpts from 300 popular pieces of music (e.g., pop songs, movie themes). For the NTKT task, we recorded simple piano melodies of folk songs using Reason. We then transposed and exported each folk melody in every key. The explicit absolute pitch training and testing programs were run on participants’ personal computers using Open Sesame software ^15^. All stimuli and training scripts are available on the Open Science Framework.

#### 2.2.2 Tests of Absolute Pitch

We assessed AP ability before and after the training program using several tests of AP. One of these tests (hereafter referred to as the UCSF Test) has been used in several prior studies of AP ^16,17^, and could be accessed through the University of California San Francisco AP website. We recorded the audio stimuli (40 piano tones, 40 sine wave tones) from the website and administered the test offline and in the lab to monitor the participants (minimizing the possibility of using non-absolute cues or strategies such as humming to artificially inflate their scores). We removed the four highest tones for the sine wave test and the four lowest tones for the piano test, which is standard for scoring the UCSF Tests. Participants identified each note in writing.

Both the UCSF Piano and Sine Tests consisted of 40 trials. Each tone lasted 1000ms with approximately a 2250ms interlude between tone onsets (meaning the inter-note-interval was approximately 3250ms). There was a longer break (10s) after every 10 notes, which gave participants the opportunity to check to make sure that they were categorizing the correct trial number. Under no circumstances were participants allowed to repeat any of the notes. Participants were randomly assigned to take either the Piano or Sine Test first.

The second AP test (hereafter referred to as the UCSD Test) was taken from Diana Deutsch’s website (deutsch.ucsd.edu), and has been used to assess AP ability across a wide variety of participants ^18^. To minimize the use of relative pitch as a cue, all intervals between successive notes were larger than an octave. We downloaded the audio file from the website and administered the test offline and in the lab to monitor the participants (minimizing the possibility of using non-absolute cues or strategies such as humming to artificially inflate their scores). The participants identified each note in writing.

The UCSD Test consisted of 36 scored piano notes, which spanned from C [3] (below middle C) to B [5] (almost three octaves above middle C) divided into three block of 12 notes. The first four notes were not scored and were designated as practice trials, which is standard for this test. Each piano note lasted approximately 500ms, and there was approximately 3750ms of silence between notes (meaning the inter-note-interval was approximately 4250ms). After each block of 12 notes, there was an extended break of approximately 18s, which gave participants the opportunity to check to make sure that they were categorizing the correct trial number. Under no circumstances were participants allowed to repeat any of the notes. The AP test we developed for the purposes of this study (hereafter referred to as the Chicago Test) was identical to the WT from training. We treated the Piano Block separately from the Multiple Timbre Block. All response time analyses associated with the Chicago Test remove outliers (greater than three standard deviations above the mean response time). The outlier cutoffs were separately calculated for each session.

#### 2.2.3 Tests of Auditory Working Memory

The implicit note memory (INM) task has been previously used as a test of auditory working memory precision^11^, and has also been associated with both explicit and implicit AP representations^11,19^. On each trial, participants heard a brief (200ms) sine wave target note, which was then masked by 1000ms of noise. Participants then had to adjust a starting note, by clicking on upward and downward arrows on the computer screen, to try to recreate the originally heard target note. The arrows moved the pitch either 10 or 20 cents up or down, depending on whether participants were clicking on the smaller arrows (10 cents) or larger arrows (20 cents). When participants believed that they had successfully recreated the original target note, they pressed a key to move onto the next trial.

There were a total of 27 sine waves in the distribution. The lowest frequency was 471.58 Hz, corresponding to a 20-cent sharp A# [4] and the highest frequency was 547.99 Hz, corresponding to a 20-cent flat C# [5]. The intermediary frequencies were evenly spaced in 10cent increments. On each trial, participants had to recreate one of 10 possible targets (corresponding to pitches 5, 7, 9, 11, 13, 15, 17, 19, 21, or 23 in the distribution), starting from one of four fixed locations (corresponding to pitches 1, 3, 25, or 27 in the distribution). Participants randomly heard all combinations of target / starting location twice, resulting in 80 trials (10 target notes × 4 starting notes × 2 repetitions). The INM task was run using the Psychophysics Toolbox in Matlab. ^20,21^

The auditory n-back (NB) task required participant to monitor the identity of spoken letter strings and remember whether the current spoken letter matched the letter presented *n* trials previously. All participants completed a 2-back and a 3-back task. For all the trials in which the current letter did not match the spoken letter presented n trials previously, participants were instructed to press a button labeled “Not Target.” Both the 2-back and 3-back consisted of 90 trials (three runs of 30 spoken letters). Letters were spoken sequentially, with an inter-stimulus-interval of 3000ms. Targets occurred one-third of the time, while non-targets occurred two-thirds of the time. Before the 2-back and 3-back, participants completed a practice round of 30 trials to familiarize themselves with the task. The NB was run in E-Prime 2.0 (Psychology Software Tools: Sharpsburg, PA).

We assessed auditory short-term memory (STM) using the auditory digit span (ADS) task. For the ADS task, participants initially heard three trials of five-number strings (e.g., 27483). There was 1000ms of silence between spoken numbers. Participants needed to correctly identify a majority (at least two) of the five-number strings in order to advance to six-number strings. If participants could not correctly identify a majority of the five-number strings (attaining zero or one correct answer), they were given three trials of four-number strings. This process of adding or removing a number based on performance repeated eight times. Thus, a perfect performance would yield a digit span score of 13 (5+8), while a completely inaccurate performance would yield a digit span score of 1. The ADS was run in E-Prime 2.0.

#### 2.2.4 Questionnaires

Participants filled out a music experience questionnaire at the completion of training, which asked about primary music experience, the number of instruments played (as well as the number of years of active musical instruction on each instrument), and the age of beginning musical instruction. Additionally, participants filled out a follow-up questionnaire, which specifically asked about participants’ explicit AP training in the time between posttest and follow-up.

### 2.3 Procedure

After providing informed written consent, participants completed the NB, INM, and ADS measures, which were followed by the AP Tests (UCSF and Chicago). These were completed in the lab over in a single session. Participants were then given portable drives containing the AP Training Program, which contained detailed instructions for completion. Participants were additionally walked through the general training protocol by the experimenter.

Over the next eight weeks, participants completed both phases of their training program as specified by the instructions. One participant (S5) was unable to complete three days of training in Week 7. Every Friday, participants completed their weekly tests. Participants uploaded their data to a secure server after each week.

Within one week of completing the eight-week training program, participants returned to the lab to complete the external AP tests (UCSF and UCSD), in addition to a musical experience questionnaire. We treated participants’ final WT as their Chicago posttest, as this was completed on the final day of the training program. The follow-up test, which was identical in design to the posttest, was conducted approximately four months after training had ended depending on availability of the participants. No feedback was given at any point during testing. Around the same time as the follow-up test, participants completed an online questionnaire that assessed their musical activities in the period between the posttest and follow-up, including explicit AP practice.

## 3. Results

### 3.1 Pre-Training

#### 3.1.1 WM and STM Assessments

Results from the pretest session confirmed that all participants scored well on the measures of auditory WM and STM. The average auditory n-back (NB) scores (using d-prime measurements ^22^) were 4.05 (*SE*: 0.17) for the 2-back, and 3.52 (*SE*: 0.32) for the 3-back. For reference, in a previous study ^11^ the average auditory 2-back score was 3.37 (*SE:* 0.16) and the average 3-back score of 2.27 (*SE:* 0.19) from a sample primarily consisting of University of Chicago undergraduates. The average error in the implicit note memory (INM) task was 2.75 (*SE*: 0.30) steps (corresponding to 27.5 cents) meaning that participants were able to recreate a briefly presented sine wave tone within approximately one-quarter of one semitone. For reference, the error of genuine AP possessors performing the identical task is on average 2.44 (*SE:* 0.19) steps ^23^. In an auditory digit span (ADS) task, participants correctly recalled an average of 8.83 (SE: 0.65) spoken digits, which was between the range of non-AP musical controls (8.1) and genuine AP possessors (10.0) previously reported ^24^.

#### 3.1.2 AP Assessments

Prior to training, no participant reached or exceeded the AP cutoffs that have been previously established in the external UCSF AP Test. ^17^ We scored the UCSF Test by giving full credit for a correct answer and three-quarters credit for answers that fell within one semitone of the correct answer, as this scoring rubric has been previously adopted for this test.^16^ For all reported AP tests, we also calculated participants’ mean absolute deviation (MAD) from the correct note in semitones, with scores of 0 reflecting perfect performance (i.e. 0 semitones removed from the correct note) and scores of 3 reflecting random guessing (i.e. uniform distribution of errors ranging from 0 to 6 semitones removed). Out of a maximum score of 36, prior to training, participants averaged 8.92 points in the Sine Test (MAD: 2.79, *SE*: 0.24) and 10.38 points in the Piano Test (MAD: 2.62, *SE*: 0.29). This level of performance was below the cutoff for the highest designation of AP (*AP*-1), defined as a minimum score of 24.5 on the Sine Test, as well as below the cutoff for the lowest AP designation (*AP-4*), defined as a minimum score of 27.5 on the Piano Test, as specified by the creators of the test.^17^ Moreover, all participants scored within the range of non-AP participants from prior investigations using this test. The distribution of pre-training responses for each participant is shown in Figure 1A (Piano) and Figure 1B (Sine), with average MAD represented in Figure 1D. All participant scores (out of 36) are reported in Table 1.

**Table 1:** Performance measures across the Chicago Piano (C-P) and Multiple Timbre (C-MT) Blocks, the UCSF Piano (UCSF-P) and Sine (UCSF-S) Tests, as well as the UCSD Test.

| *Pretest*<br>Participant | C-P | C-MT | UCSF-P | UCSF-S | UCSD |
| --- | --- | --- | --- | --- | --- |
| S1 | 25.00% (7.14s) | 25.00% (4.08s) | 9.25 | 3 | NA |
| S2 | 62.50% (2.81s) | 52.08% (3.19s) | 16.25 | 12.75 | NA |
| S3 | 14.58% (4.13s) | 12.50% (4.36s) | 5.75 | 7.5 | NA |
| S4 | 20.83% (7.71s) | 12.50% (7.08s) | 8.5 | 8.5 | NA |
| S5 | 70.83% (3.31s) | 68.75% (2.90s) | 15.5 | 13.5 | NA |
| S6 | 6.25% (6.90s) | 4.17% (6.44s) | 7 | 8.25 | NA |
| *Posttest*<br>Participant | C-P | C-MT | UCSF-P | UCSF-S | UCSD |
| S1 | 52.08% (7.08s) | 54.17% (6.25s) | 13.25 | 4.75 | 41.67% |
| S2 | 97.92% (1.53s) | 100% (1.48s) | 34.5 | 29.5 | 94.44% |
| S3 | 37.50% (2.43s) | 31.25% (2.16s) | 10.5 | 11.5 | 19.44% |
| S4 | 47.92% (4.48s) | 52.08% (5.51s) | 7.5 | 6 | 11.11% |
| S5 | 100% (2.81s) | 100% (2.79s) | 24.75 | 17.75 | 97.22% |
| S6 | 0% (2.32s) | 0% (2.58s) | 16.75 | 5.5 | 2.78% |
| *Follow-up*<br>Participant | C-P | C-MT | UCSF-P | UCSF-S | UCSD |
| S1 | 56.25% (6.48s) | 52.08% (5.84s) | 13.5 | 6.25 | 63.89% |
| S2 | 93.75% (1.73s) | 91.67% (2.02s) | 30.75 | 32.75 | 77.78% |
| S3 | 33.33% (2.65s) | 47.92% (2.34s) | 9.5 | 7 | 52.78% |
| S4 | 18.75% (6.21s) | 22.92% (6.34s) | 11 | 7 | 19.44% |
| S5 | 93.75% (3.01s) | 91.67% (2.81s) | 24.75 | 18.25 | 91.67% |
| S6 | 2.08% (2.92s) | 0% (3.07s) | 13.25 | 5 | 0% |

*Note:* Parentheses following the C-P and C-MT percentages represent mean response time. UCSF values represent performance (out of 36 scored trials), where full credit was given for correct answers and three-quarters credit was given for incorrect answers within one semitone of the correct answer. UCSD values represent percentage correct (no credit for semitone errors).

**Figure 1:**
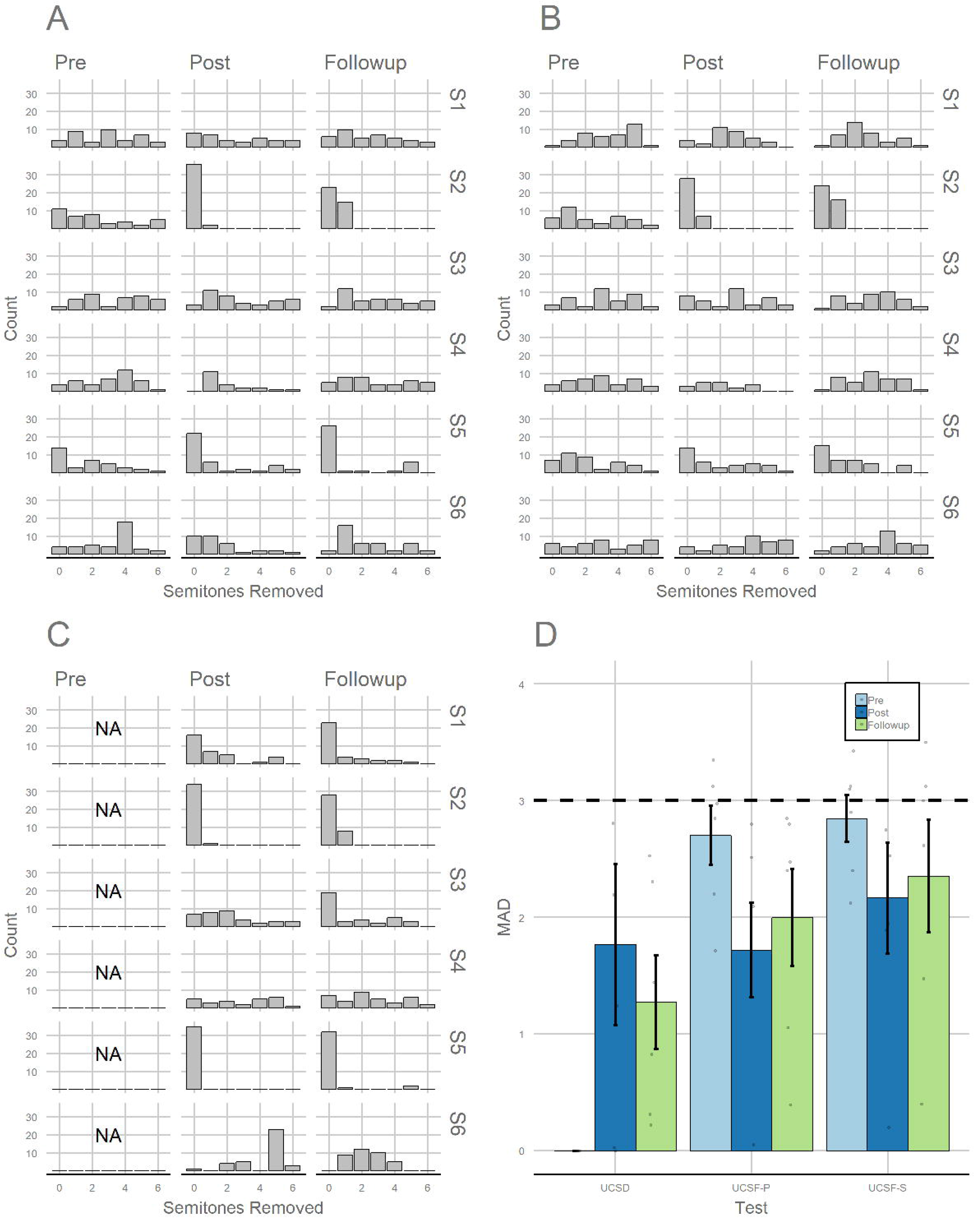
Distribution of semitone errors for each participant (S1-S6) and each testing session (Pre, Post, and Follow-Up) for the UCSF Piano Test (A), UCSF Sine Test (B), UCSD Test (C), and mean absolute deviation across all participants for these tests (D).

Despite not granting credit for semitone errors on the Chicago Test, we observed higher performance compared to the UCSF Test, likely because the Chicago Test was self-paced. Participants achieved 33.3% accuracy for the Piano Block (MAD: 1.96, *SE*: 0.45) and 29.2% accuracy for the Multiple Timbre Block (MAD: 1.98, *SE*: 0.44). Participants’ mean response times (RTs) were 5.33s (*SE*: 0.88s) for the Piano Block and 4.67s (*SE*: 0.70s) for the Multiple Timbre Block, which was slower than the fixed presentation rate of notes in the UCSF Test. The distribution of responses for each participant are represented Figure 2A (Piano Block) and Figure 2B (Multiple Timbre Block), with mean accuracy represented in Figure 2C and mean response time represented in Figure 2D. All participants’ accuracy and RTs are additionally reported in Table 1.

**Figure 2:**
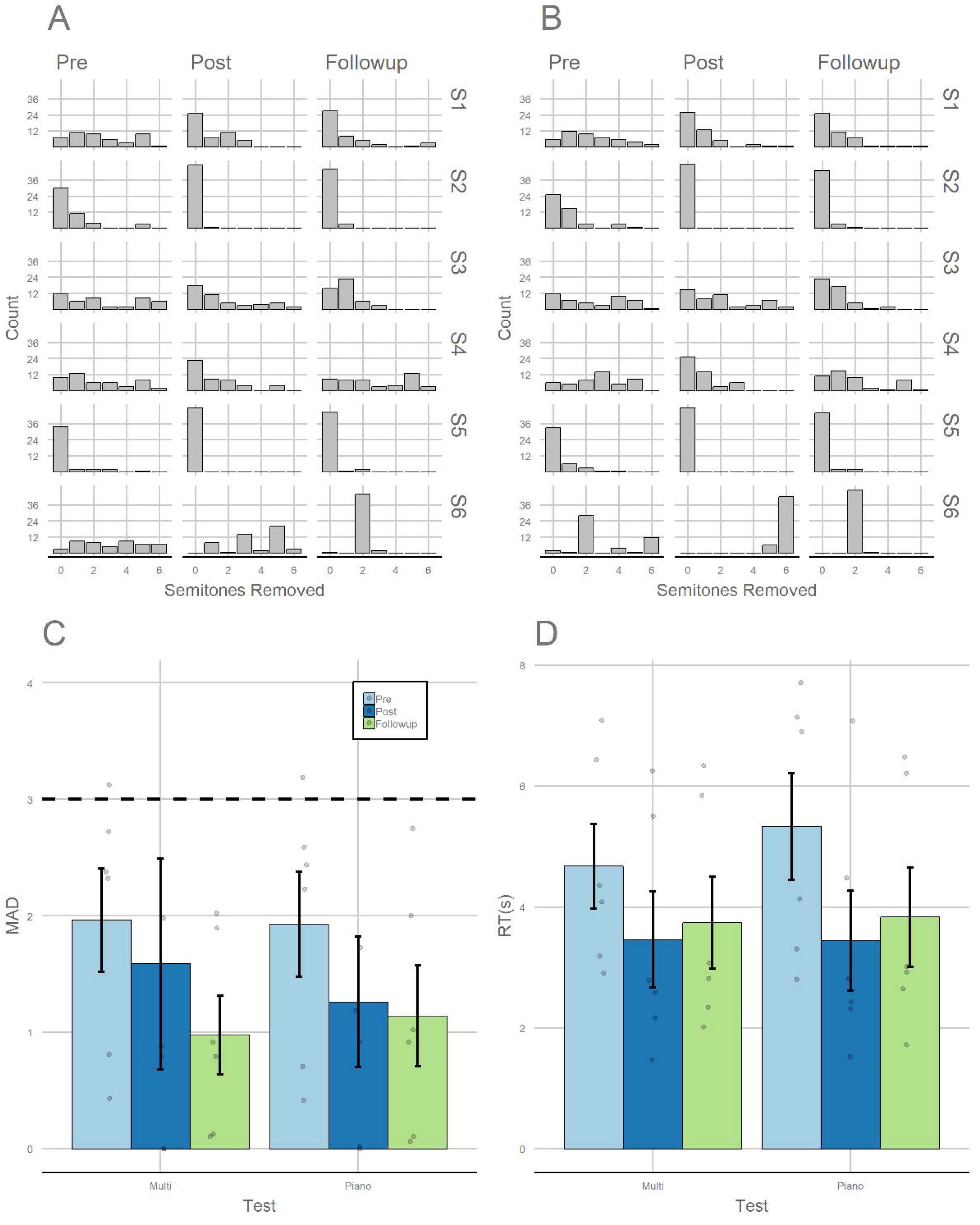
Distribution of semitone errors for each participant (S1-S6) and each testing session (Pre, Post, and Follow-Up) for the Chicago Test, split by the Piano Block (A) and Multiple Timbre Block (B). Mean absolute deviation across all participants for this test is represented in panel C, whereas mean response time across all participants for this test is represented in panel D.

### 3.2 Training

The complete training data are available on Open Science Framework. Here, we focus on performance from the end-of-week tests (WT for First and Second Phases, NTKT for Second Phase).

#### 3.2.1 Isolated Note Classification

Participants displayed an improvement in classifying isolated notes without feedback over the course of training. After the first week of training, participants correctly categorized 46.2% of notes for the Piano Block (*MAD*: 1.33, *SE*: 0.40) and 49.0% of notes for the Multiple Timbres Block (*MAD*: 1.43, *SE*: 0.50), which already represented an improvement from pre-training performance. By the end of the seventh week of training, performance had increased to 55.2% of notes for the Piano Block (*MAD*: 1.02, *SE*: 0.52) and 55.6% of notes for the Multiple Timbres Block (*MAD*: 1.07, *SE*: 0.52). In terms of response time, participants were nominally *slower* in classifying notes compared to the pre-training assessment after the first week of training.

Participants took 5.78s (*SE:* 0.87s) to respond to notes in the Piano Block and took 5.42s (*SE:* 0.71s) to respond to notes in the Multiple Timbres Block. By the end of the seventh week, however, participants were responding more quickly on average, taking 3.90s (*SE:* 0.86s) to respond to notes in the Piano Block and 4.09s (*SE*: 0.93s) to respond to notes in the Multiple Timbres Block. Figure 3A plots MAD as a function of training week, while Figure 3B plots response time as a function of training week.

**Figure 3:**
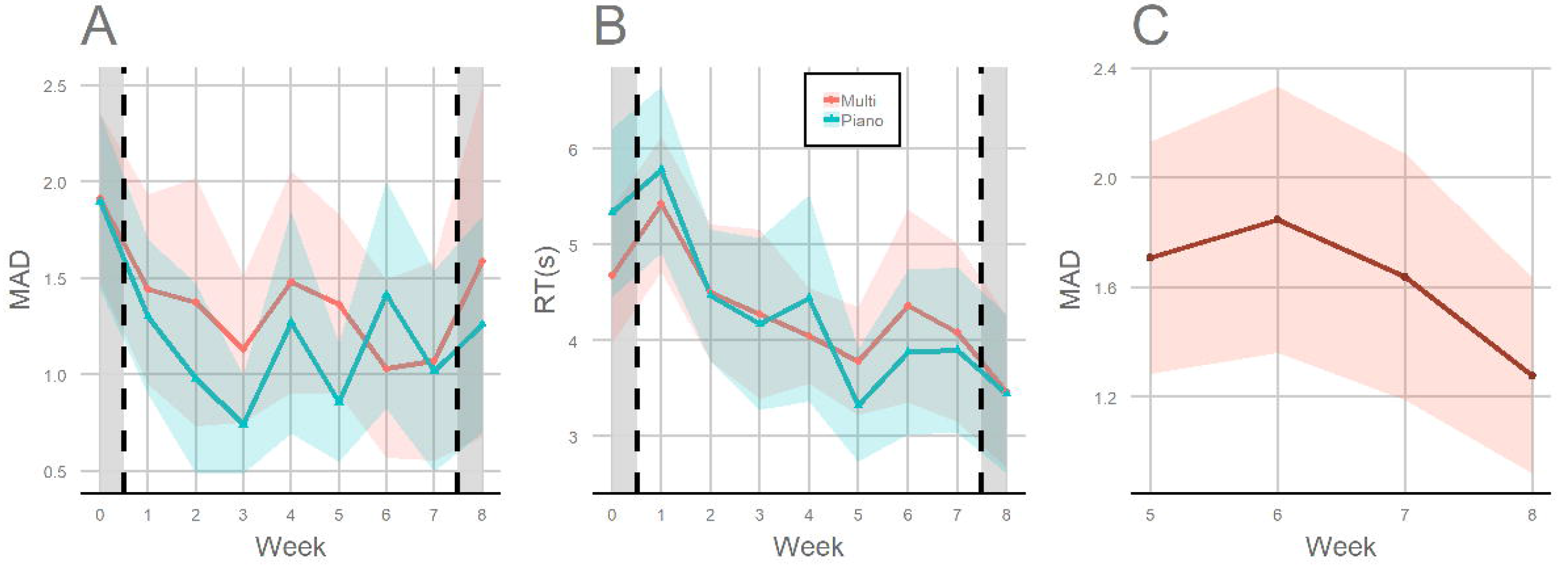
Averaged results from the weekly assessments during training. The weekly test (WT) assessment is separated by mean absolute deviation (A) and response time (B). The name that key test (NTKT) is represented in terms of mean absolute deviation.

#### 3.2.2 Key Signature Classification

Despite only being administered four times (with each administration only consisting of 12 trials), we observed improvements in key signature classification over the course of training. In the first administration of the NTKT, participants correctly identified the key signature on 34.7% of trials (*MAD:* 1.71, *SE*: 0.42). By the final administration of the NTKT, participants correctly identified the key signature on 41.7% of trials (*MAD*: 1.28, *SE*: 0.36). Figure 3C plots MAD for the NTKT as a function of week.

### 3.3 Post-Training

#### 3.3.1 Immediate Test

Within one week after training had ended, we retested all participants inside the lab on the UCSF Test and an AP test not administered prior to training (the UCSD Test) ^18^. Participant S2 showed considerable improvement on the UCSF Piano and Sine Tests, scoring 34.5 on the Piano Test (MAD: 0.05) and 29.5 on the Sine Test (MAD: 0.20) - a level that was above the cutoff for *AP-1* ability. Participant S5 also showed substantial improvements, scoring 24.75 on the Piano Test (MAD: 1.32) and 17.75 on the Sine Test (MAD: 1.89), though participant S5 missed the *AP-1* qualification by 6.75 points and missed the *AP-4* qualification by 2.75 points. All other participants missed the cutoff for *AP-1* by at least 13 points and the cutoff for *AP-4* by at least 10.75 points (Table 1). On the UCSD Test, Participant S2 scored 94.44% (MAD: 0.03), while Participant S5 scored 97.22% (MAD: 0).^1^ This level of performance on the UCSD Test qualifies both participants as AP possessors based on previous interpretations of this test (using an 85% accuracy cutoff for conservative AP inclusion) ^18^. As a comparison, the third highest scoring participant (S1) achieved 44.4% accuracy (MAD: 1.24), which was above chance by over 36 percentage points (represented by 1/12, or 8.33%) but below typical thresholds used to identify AP. The distribution of responses for each participant on the UCSD Test are represented Figure 1C, with average MAD represented in Figure 1D.

This pattern of results extended to the Chicago Test as well, with participants S2 and S5 displaying virtually perfect performance on both the Piano and Multiple Timbre Blocks. Participant S2 achieved 97.9% accuracy on the Piano Block (MAD: 0.02) and 100% on the Multiple Timbre Block (MAD: 0), while participant S5 achieved 100% on both the Piano Block and Multiple Timbre Block (MAD: 0). For reference, the third highest performing participant (S1) scored 52.1% on the Piano Block (MAD: 0.92) and 54.2% on the Multiple Timbre Block (MAD: 0.88), which once again demonstrates clear above-chance performance but does not reach thresholds typically used to denote genuine AP.

#### 3.3.2 Follow-Up Test

To assess the stability of AP category learning, we retested all participants approximately four months (*M* = 128.17 days, *SD* = 6.71 days, range of 117 - 134 days) after training had ended. No participant had reported actively rehearsing pitch-label associations in the time between the immediate AP posttests and the follow-up tests. Previous AP training research has been criticized for not following up with participants after training has ended to see how category learning is retained ^25^, as genuine AP possessors appear to have relatively stable categories that do not require explicit maintenance (though see ^13,14,26^ for alternate views). We administered the same AP assessments given to participants in the posttest (UCSF Test, the UCSD Test, and the Chicago Test).

Results from the follow-up AP tests supported the conclusion that Participants S2 and S5 retained stable performance across all AP assessments. For the UCSF Test, Participant S2 still passed the *AP-1* cutoff, scoring 32.75 on the Sine Test and 30.75 on the Piano Test. Participant S5 retained high performance (13.75 on Sine, 24.75 on Piano), but was still below the *AP-1* cutoff by 10.5 points and missed the *AP-4* designation by 2.75 points. For the UCSD Test, Participant S2 scored 77.78% (MAD: 0.22) and Participant S5 scored 91.67% (MAD: 0.31).

Even though Participant S2 fell below the 85% cutoff used as a conservative inclusion measure in prior studies, S2 never missed by more than one semitone. Adopting the more liberal inclusion measures from prior studies that have used the UCSD Test (in which semitone errors are allowed), Participant S2 would still be categorized as possessing AP ^18^. It should be noted that Participant S2 also took the UCSD Test again approximately 16 months post-training for a separate study and achieved 88.89% accuracy (MAD: 0.14) not including semitone errors as correct. Finally, for the Chicago Test, Participants S2 and S5 scored identically in terms of percentage correct (93.75% on the Piano Block, MADs of 0.06 and 0.10, respectively) and 91.67% on the Multiple Timbre Block, MADs of 0.11 and 0.13, respectively).

The third highest performing participant was once again S1, who displayed clear above chance performance on the UCSD and Chicago Tests but did not reach thresholds typically used to define AP. Participant S1 scored 63.9% on the UCSD Test (75% when including semitone errors as correct), 56.3% (MAD: 1.02) on the Piano Block of the Chicago Test, and 52.1% (MAD: 0.91) on the Multiple Timbres Block of the Chicago Test.

### 3.4 Comparisons with External Studies

One strength of the present experiment is that AP performance was assessed with three separate tests. Two of these tests - the UCSF Test and the UCSD Test - have been externally administered in a number of previous studies ^17,27–31^, which makes their interpretation (i.e. in terms of AP thresholds) more straightforward. Yet, we acknowledge that these two tests in particular also may reduce the dimensionality of AP by adopting a fixed note presentation rate. This is because individuals who are able to keep up with the presentation rate will cluster together as an “AP group” (regardless of individual variation in speed) and individuals who are not able to keep up with the presentation rate will cluster together as a “non-AP group” (regardless of individual variation in speed). As such, individuals who may exhibit some intermediary AP ability will be pulled toward one of these two groups depending on whether their classification speed is sufficient to keep up with the particular test, possibly exaggerating the dichotomous nature of AP.

In this sense, our untimed Chicago Test provides a richer means of capturing gradations in AP ability (including at the top end of the performance spectrum), especially if speed and accuracy are jointly considered. To directly test whether a simultaneous consideration of speed and accuracy supports or challenges the observation that Participants S2 and S5 were behaviorally indistinguishable from genuine AP possessors post-training, we directly compare the data from the present experiment with an influential prior investigation of AP (n = 51), hereafter referred to as the “McGill Test” ^12^. The McGill Test is particularly well-suited for comparisons with the Chicago Test because (1) both were administered on the computer, (2) both required participants to click on one of twelve provided note categories, arranged in a circular fashion, (3) both did not adopt a “timeout” window or present notes at a fixed rate, and (4) both have associated data that span a full range of performance profiles (spanning from perfect to chance performance). These parallels make it possible to better interpret the results from the present experiment in a broader context treating AP as a more distributed ability, as well as better assess whether our highest-performing participants were comparable to the highest-performing AP participants in this prior investigation.

To facilitate the interpretation between the Chicago Test and the McGill Test, we created an index that incorporated both MAD and log response time (logRT), such that slower RTs would be penalized relative to faster RTs. This index, which was specifically created by adding 10 to an individual’s MAD and then multiplying this number by their logRT, was previously shown by the authors of the McGill Test to be sensitive to gradations in AP ability, capturing the nominal categories of “AP” (near perfect and fast), “non-AP” (near random and slow), and a wide range of intermediate AP abilities ^12^.

To measure how the participants in the present experiment compared to the participants from this previous dataset, we adopted the following procedure. First, we extracted the McGill Test data by digitizing the scatterplot (represented as Figure 7 in their paper). This provided us with all 51 participants’ index values as well as their overall accuracy (0-100%). Second, we ran a k-means clustering algorithm on these data to define three groups (nominally: *genuine AP*, *pseudo AP*, and *non AP*).^2^ Third, we used a linear discriminant analysis to classify participants into one of these three groups based on their index value and mean accuracy. Fourth, we used the results of this linear discriminant analysis to predict the classification of the six participants in the present experiment at each time point (pre-training, immediate posttest, follow-up). In addition to receiving a nominal classification (i.e. non AP, pseudo AP, or genuine AP), this analysis also provided the posterior probability of belonging to each category.

Figure 4 (left column) plots the index value (logRT and MAD) against note classification accuracy, overlaying the six participants from the present experiment onto the extracted McGill Test data. Prior to training (“PRE”), Participant S2 and S5 were clearly distinguishable from the other four participants in terms of both the index value and mean accuracy. Yet, the classifier placed both participants into the intermediate (“pseudo-AP”) group (100% posterior probability for S2, 98.4% posterior probability for S5). All other participants were classified in the lowest (“non-AP”) group (100% posterior probability for S1, S4, and S6, 99.9% posterior probability for S3).

In the immediate posttest (“POST”), Participants S2 and S5 were both classified as belonging to the highest (“genuine”) AP Group (100% posterior probability). Participants S1, S3, and S4 were all classified as belonging to the pseudo-AP group (100%, 96.3%, and 100% posterior probabilities, respectively), and Participant S6 was classified in the non-AP group (100% posterior probability). These results were generally consistent in the follow-up test (“FOLLOW”), with the exception of Participant S4 whose performance sufficiently worsened to be classified in the non-AP group. Participants S2 and S5 were still classified as belonging to the genuine-AP group, Participants S1 and S3 were still classified as belonging to the pseudo-AP group, and Participant S6 was still classified as belonging to the non-AP group (all 100% posterior probabilities). Overall, these analyses suggest that Participants S2 and S5 were performing sufficiently well post-training to be classified within the highest AP Group, *even though their performance prior to training was distinguishably lower than the highest, genuine-AP group.*

**Figure 4:**
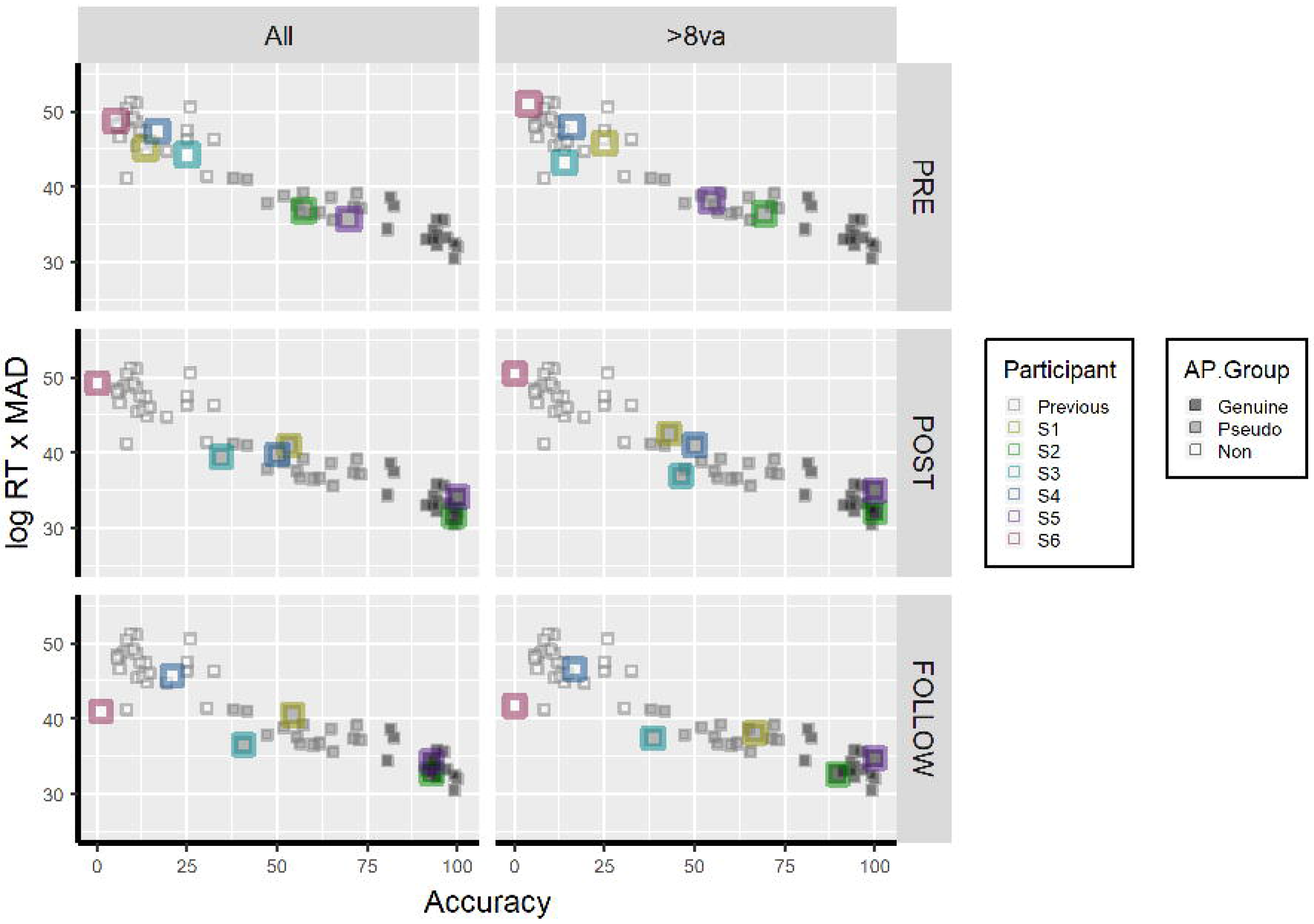
Comparison of the present experiment with Bermudez and Zatorre (2009), separated by testing session (Pre, Post, and Follow-Up). The left column (“All”) represents performance across all trials, while the right column (“>8va”) represents performance on trials in which there was more than an octave separating the heard note from the previous note. An index consisting of mean absolute deviation and log response time is represented on the y-axis, while percent correct is represented on the x-axis. Prior to training, no participant was classified in the highest (“genuine”) AP group for either analysis. In the immediate test post-training, as well as in the follow-up test four months after training, participants S2 and S5 performed indistinguishably from the highest AP performers from Bermudez and Zatorre (2009) in both analyses.

### 3.5 Examining Alternative Strategies

An alternative explanation of the present results is that participants did not actually improve the resolution of their absolute note categories, but rather learned one or two notes and then used relative pitch to infer the rest. This kind of strategy requires some AP memory (and is thus of interest to understanding AP as a distributed ability), but it has been considered as distinct from genuine AP from the earliest scientific investigations of absolute pitch memory ^32^. To assess the evidence supporting the possibility that participants had simply memorized one or two notes, we carried out an analysis that was conceptually identical to the McGill Test analysis described in the previous section. The main difference, however, was that we excluded trials in which the note was separated from the preceding trial’s note by one octave or less. The reasoning behind this exclusion is that shifts in octave are thought to disrupt individuals using a relative pitch strategy, and several AP tests-including the UCSD Test and the McGill Test - change octaves on each trial for this reason. While increased response times for these “mixed octave” trials would not necessarily be surprising, as even genuine AP possessors demonstrate increased response time to identifying notes that vary across octaves ^33^, the critical question is whether the interpretation of participants’ AP abilities before and after training is meaningfully affected by this restricted and more conservative analysis.

The results are plotted in Figure 4 (left column). Prior to training, Participants S2 and S5 could still be differentiated from the other participants; however, they were classified in the pseudo-AP group (99.3% posterior probability for S2, 100% posterior probability for S5). Interestingly, Participant S2 was closer to the genuine-AP group than Participant S5, which represented a reversal of the full trial analysis. This suggests that Participant S2 performed better on these mixed octave trials, whereas Participant S5 performed worse. All other participants were classified in the non-AP group (100% posterior probability).

In the immediate posttest, Participants S2 and S5 were classified in the genuine-AP group (100% posterior probability). Participants S1, S3, and S4 were all classified in the pseudo-AP group, though the rankings of these participants differed from the full trial analysis. Specifically, Participant S1 - who was the most accurate pseudo-AP participant when considering all trials - was the *least* accurate pseudo-AP participant in this analysis (99.1% posterior probability for pseudo-AP, 0.9% for non-AP). This suggests that Participant S1 may have been using a strategy in the immediate posttest that was particularly harmed by octave changes. In the follow-up test, however, Participant S1 displayed improvements in accuracy (42.9% to 66.7%), MAD (1.21 to 0.63), and response time (7.19s to 5.08s) and was thus the highest-performing pseudo-AP participant (100% posterior probability of pseudo-AP), suggesting a possible shift in strategy that more closely reflected what is observed within a genuine AP population. Participants S2 and S5 were still classified in the genuine-AP group (100% posterior probabilities). Participant S3 was classified in the pseudo-AP group (99.9% posterior probability), and both Participants S4 and S6 were classified in the non-AP group (100% posterior probability).

Taken together, these comparisons with the McGill Test data highlight the importance of jointly using response time and accuracy to gain a more comprehensive understanding of variability in AP performance. In particular, the second analysis (examining trials that were at more than one octave removed from the previous trial) suggests that Participant S1 was achieving an intermediate level of AP performance in the immediate posttest through use of an alternate strategy, such as using one or two memorized notes to infer the rest. Participants S2 and S5, in contrast, displayed performance profiles that were precisely what would be expected from a genuine AP possessor (suggesting that training improved the resolution of all note categories, not just one or two).

## 4. Discussion

The present results challenge two assumptions regarding AP - (1) that it is a dichotomous, “all or none” ability, and (2) that genuine AP patterns of performance cannot emerge in post-critical period adults ^1^. We interpret the present experiment as providing strong evidence for adult AP acquisition; however, we acknowledge that this is not the only interpretative framework that may explain these data. Below, we outline two “non-learning” alternative explanations and discuss why they are insufficient in explaining our results.

The first non-learning explanation is that our successful participants were always “AP possessors,” and therefore the training program merely revealed an inherent ability rather than reflected genuine learning. This kind of explanation comes from a broader framework treating AP as an innate perceptual ability, requiring essentially no environmental input and therefore not being restricted to a critical period of development ^34–36^. At first glance, this possibility seems to be partially supported by the pretest results, as Participants S2 and S5 were already distinguishable from the other participants in terms of AP performance. However, the notion that both were already “AP possessors” (as typically defined) seems unlikely for two reasons. First, the data-driven approach to defining the highest cluster of AP performance (“genuine AP”) demonstrated that, prior to training, Participants S2 and S5 were not sufficiently fast and accurate to be considered a part of this group (0% probability for Participant S2, 1.6% probability for Participant S5). Second, both participants had extensive musical backgrounds, and thus had extensive opportunities to learn note-label associations. Specifically, Participant S2 and S5 began musical instruction at 7 and 6 years old, respectively, and played their primary instrument for 8 and 20 years, respectively. If AP reflects an innate ability that only nominally requires environmental shaping (i.e. to learn the conventional names of Western musical notes), then both participants should have performed with sufficient speed and accuracy to be labeled as AP possessors prior to training, given their considerable prior experience associating pitches with their respective note names.

The second non-learning explanation is that our successful participants never possessed AP (even after training), but rather found some means of augmenting performance through strategies that did not involve the actual refinement of the twelve note categories. In particular, participants may have been able to memorize one or two absolute pitches and then used relative pitch to categorize the remaining pitches - a phenomenon that has been discussed as distinct from genuine AP for the better part of a century ^32^. Yet, given the history of this distinction in the literature, many AP assessments have adopted methodologies to minimize relative pitch strategies, primarily related to note classification speed and the intervallic distance between consecutive notes. Yet, these factors did not appear to hinder our highest-scoring participants’ performance post-training. Even when using a more nuanced approach (simultaneously weighing accuracy and response speed for the untimed Chicago Test), which is more sensitive to graded AP performance profiles, we found strong evidence that Participants S2 and S5 were indistinguishable from a prior group of “genuine” AP possessors post-training, even when limiting the analyses to notes that were separated by more than one octave. Thus, if one wants to claim that what we observed is not genuine AP, then either the current definition of AP or the ways in which AP is tested need to be fundamentally reconsidered.

### 4.1 Implications for Theories of AP Acquisition

The present results clearly support a *skill acquisition* theory of AP, in that some individuals can improve their AP abilities following explicit perceptual training to the point where their performance is indistinguishable from AP possessors whose abilities manifested early in life. Yet, it is misleading to think that a *skill acquisition* theory of AP cannot be partly reconciled with more dominant theories of AP acquisition. In the following paragraphs, we highlight how the results from the present experiment may be integrated with both the *critical period* and *innate* theories of AP acquisition, particularly when conceptualizing AP as a distributed and non-dichotomous ability.

Under the *critical period* theory, AP acquisition is almost exclusively confined to an early window of development. The “cutoff’ for being able to acquire AP is not likely a strict age, but rather reflected as a decreasing probability of acquisition as a function of aging ^5^. Thus, finding two successful adult AP learners is not technically incompatible with the *critical period* theory; however, observing a 33% success rate among an adult sample would be virtually impossible given (1) the presumed rarity of AP and (2) the relative probability of acquiring AP as an adult suggested by a *critical period* framework. An important point to consider in the context of the present experiment, however, is that both successful participants began musical training at an age that would be more compatible with the *critical period* theory (i.e. younger than 8 years old). Given that both participants had experience associating musical pitches with their note labels relatively early in life, the *critical period* theory could be integrated with a *skill acquisition* theory. For example, early musical training may be necessary (but not sufficient) for developing the capacity to meaningfully refine AP ability, though the actual refinement of note categories may take on a more heterogeneous trajectory. On one end of this continuum would be individuals who acquire AP rapidly and without any reported effort ^37^ On the other end of this continuum would be individuals who refine their note categories over the course of extended, concentrated practice (as was the case in the present experiment). Importantly, just because the experiences in acquiring AP differ across this continuum does not mean that individuals cannot converge on the same level of AP proficiency.

Yet, the *skill acquisition* theory of AP does not necessarily suggest that any individual can develop sufficient speed and accuracy to be indistinguishable from a genuine AP possessor. In our own sample, the two successful AP learners were performing better *before training* than the third highest-performing participant performed *after training*. Moreover, the third-highest performing participant was slower and more disrupted by shifts larger than an octave compared to the two best learners, suggesting that performance may have been the result of adopting a different strategy compared to the two highest performers. As such, our results may also inform the *innate* theory of AP acquisition, for it is possible that only some adults can develop genuine AP levels of performance post-training. If this is the case, then it would be of great scientific interest to understand (1) the base rate of trainable adults, (2) the perceptual, cognitive, and possibly underlying genetic factors that differentiate these individuals from non-trainable adults, and (3) whether these individuals always report early musical training.

### 4.2 AP as an Auditory Skill

When framing AP as a clearly defined, dichotomous ability ^29^, it is difficult to entertain the idea that AP can be acquired as a function of training. This is because there is no middle ground - performance should be close to random or close to perfect. As such, the transition from “non-AP” to “AP” would represent a transition from the complete absence of an ability to the total manifestation of an ability, as if AP reflected a sudden insight into how to interpret auditory pitch. In contrast, when framing AP as a distributed, multidimensional ability ^U2^, learning theories become more theoretically plausible. This is because “successful” AP learning does not need to represent a binary switch. Rather, in some cases, a rather modest degree of learning may be sufficient to result in “genuine AP” levels of performance. Yet, approaching AP as a learnable skill has been largely dismissed in the literature, despite a growing body of research suggesting that, even among AP possessors, absolute pitch ability can be significantly strengthened or even weakened by environmental experiences (e.g., recent musical training) outside of a critical period ^14,38,39^.

Why, then, is there a hesitancy to consider AP in a *skill acquisition* framework? One reason is that there still may be an implicit assumption that AP should still be dichotomized, even if performance lies along a continuum. In other words, there is still a tendency to treat individuals who fall on the highest end of an AP continuum as engaging in a fundamentally different process from other individuals (e.g., differentiating “genuine” from “pseudo” AP as a character trait). While this approach may be justified in some cases (e.g., to distinguish alternative strategies in note identification), it also may unnecessarily simplify AP and downplay the role of learning and plasticity in AP more generally. A second reason why there may be hesitancy in accepting a *skill acquisition* framework is the lack of conclusive previous empirical evidence that AP can trained in adults ^37^ Of course, inferring that AP cannot be trained in adults based on the absence of prior evidence represents an acceptance of a null hypothesis, and as such it is impossible to know whether failures of previous training studies were due to participant selection, the length or nature of the training regime, the operationalization of AP ability, or other factors.

Indeed, the positive results obtained in the present experiment point may be partly attributed to the selection of participants based on exceptional auditory memory abilities. Recent research has associated aspects of auditory WM and STM with pitch memory performance in a wide variety of settings, such as (1) absolute memory for familiar musical recordings, (2) the rapid and explicit training of AP among previously naïve adults, and (3) MAD among a “genuine AP” population for perceptually challenging notes. Moreover, “genuine AP” possessors appear to have an enhanced auditory (but not visual) digit span relative to musically matched controls ^24^, which further suggests that general auditory memory abilities may be a particularly important factor in understanding absolute pitch representations, including the explicit training of AP. As such, future investigations into adult AP learning may benefit from dissociating auditory WM/STM abilities from (early) musical training, as this could provide important insights into both the *critical period* and *skill acquisition* theories of AP.

### 4.3 Conclusion

The present results give empirical weight to the theoretical treatment of AP as an auditory skill rather than a static talent. This viewpoint is supported through providing the most conclusive demonstration to date that AP ability can be improved through training to the point of being indistinguishable from “genuine AP” within some adults. It is also important to consider that these levels of AP performance were achieved with eight weeks (approximately 32 hours) of adult training, which is far less than the amount of training that is thought to be required for the explicit learning of AP in childhood based on prior research ^40^.

While the present study is limited in sample size, the demonstration that even two adults can, with moderate training, reach genuine AP levels of performance is theoretically important. No prior published study has demonstrated a comparable level of successful adult AP learning and long-term retention. As such, we stress the importance of these findings as proof-of-concept that adult AP acquisition is possible and that learning remains stable months after explicit training has ended. To this end, *any* demonstration of successful AP learning by an adult will inform the discussion of the underlying mechanisms of AP acquisition and maintenance. While the present results cannot refute either the *critical period* or *innate* theories of AP acquisition, they suggest that aspects of both theories should be more fully integrated with a *skill acquisition* theory of AP. This integration becomes clearer when conceptualizing AP as a distributed and multifaceted ability rather than a static and dichotomous ability. Overall, it is our hope that these results will refocus future inquiry regarding AP to treat the ability as distributed and at least partly plastic, even into adulthood, as this refocusing will lead to a more complete understanding of how humans perceive and remember absolute pitch information more generally.

## Acknowledgements

*This research was supported by the Multidisciplinary University Research Initiatives (MURI) Program of the Office of Naval Research through grant, DOD/ONR N00014-13-1-0205.*

## Note on Open Materials

*All data and training programs associated with this paper are available on the Open Science Framework (https://osf.io/9n48c/)*

## Footnotes

1 The discrepancy between accuracy and MAD for participant S5 was because one note was not labeled at all. As such, it was counted as incorrect but could not be used in the calculation of MAD.

2 We acknowledge that defining three clusters may be viewed as arbitrary. However, it should be noted that doubling the number of clusters (*k*=6) does not change the interpretation of the highest-performing participants (S2 and S5).

## References

1. Takeuchi, A. H. & Hulse, S. H. Absolute pitch. Psychol. Bull. 113, 345–361 (1993).

2. Deutsch, D. Absolute Pitch. in The Psychology of Music (ed. Deutsch, D.) 141–182 (Academic Press, 2013). doi:10.1016/B978-0-12-381460-9.00005-5

3. Ward, W. D. & Burns, E. M. Absolute pitch. in The Psychology of Music (ed. Deutsch, D.) 431–451 (Academic Press, 1982).

4. Crozier, J. B. Absolute pitch: Practice makes perfect, the earlier the better. Psychol. Music 25, 110–119 (1997).

5. Levitin, D. J. & Zatorre, R. J. On the Nature of Early Music Training and Absolute Pitch: A Reply to Brown, Sachs, Cammuso, and Folstein. Music Percept. 21, 105–110 (2003).

6. Hartman, E. B. The influence of practice and pitch-distance between tones on the absolute identification of pitch. Am. J. Psychol. 67, 1–14 (1954).

7. Cuddy, L. L. Practice effects in the absolute judgment of pitch. J. Acoust. Soc. Am. 43, 1069–1076 (1968).

8. Lundin, R. W. Can perfect pitch be learned? Music Educ. J. 49, 49–51 (1963).

9. Vianello, M. a & Evans, S. H. Note on pitch discrimination learning. Percept. Mot. Skills 26, 576 (1968).

10. Gervain, J. et al. Valproate reopens critical-period learning of absolute pitch. Front. Syst. Neurosci. 7, 1–11 (2013).

11. Van Hedger, S. C., Heald, S. L. M., Koch, R. & Nusbaum, H. C. Auditory working memory predicts individual differences in absolute pitch learning. Cognition 140, 95–110 (2015).

12. Bermudez, P. & Zatorre, R. J. A distribution of absolute pitch ability as revealed by computerized testing. Music Percept. An Interdiscip. J. 27, 89–101 (2009).

13. Hedger, S. C., Heald, S. L. M. & Nusbaum, H. C. Absolute Pitch May Not Be So Absolute. Psychol. Sci. 24, 1496–1502 (2013).

14. Wilson, S. J., Lusher, D., Martin, C. L., Rayner, G. & McLachlan, N. Intersecting factors lead to absolute pitch acquisition that is maintained in a ‘Fixed do’ Environment. Music Percept. 29, 285–296 (2012).

15. Mathôt, S., Schreij, D. & Theeuwes, J. OpenSesame: An open-source, graphical experiment builder for the social sciences. Behav. Res. Methods 44, 314–324 (2012).

16. Athos, E. A. et al. Dichotomy and perceptual distortions in absolute pitch ability. Proc. Natl. Acad. Sci. U. S. A. 104, 14795–14800 (2007).

17. Baharloo, S., Johnston, P. a, Service, S. K., Gitschier, J. & Freimer, N. B. Absolute pitch: an approach for identification of genetic and nongenetic components. Am. J. Hum. Genet. 62, 224–231 (1998).

18. Deutsch, D., Henthorn, T., Marvin, E. & Xu, H. Absolute pitch among American and Chinese conservatory students: Prevalence differences, and evidence for a speech-related critical period. J. Acoust. Soc. Am. 119, 719 (2006).

19. Van Hedger, S. C., Heald, S. L. M. & Nusbaum, H. C. Long-term pitch memory for music recordings is related to auditory working memory precision. Q. J. Exp. Psychol. 1–13 (2017). doi:10.1080/17470218.2017.1307427

20. Brainard, D. H. The Psychophysics Toolbox. Spat. Vis. 10, 433–436 (1997).

21. Pelli, D. G. The VideoToolbox software for visual psychophysics: transforming numbers into movies. Spatial vision 10, 437–442 (1997).

22. Macmillan, N. A. & Creelman, C. D. Detection theory: A user’s guide. (Lawrence Erlbaum Associates, 2005).

23. Heald, S. L. M., Van Hedger, S. C. & Nusbaum, H. C. Auditory category knowledge in experts and novices. Front. Neurosci. 8, 1–15 (2014).

24. Deutsch, D. & Dooley, K. Absolute pitch is associated with a large auditory digit span: A clue to its genesis. J. Acoust. Soc. Am. 133, 1859–1861 (2013).

25. Rush, M. An experimental invesitagtion of the effectiveness of training on absolute pitch in musicians. 401 (1989).

26. Van Hedger, S. C., Heald, S. L. M., Uddin, S. & Nusbaum, H. C. A Note by Any Other Name: Intonation Context Rapidly Changes Absolute Note Judgments. J. Exp. Psychol. Hum. Percept. Perform. (2018). doi:10.1037/xhp0000536

27. Baharloo, S., Johnston, P. A., Service, S. K., Gitschier, J. & Freimer, N. B. Absolute Pitch: An Approach for Identification of Genetic and Nongenetic Components. Am. J. Hum. Genet. 62, 224–231 (1998).

28. Baharloo, S., Service, S. K., Risch, N., Gitschier, J. & Freimer, N. B. Familial aggregation of absolute pitch. Am. J. Hum. Genet. 67, 755–758 (2000).

29. Athos, E. A. et al. Dichotomy and perceptual distortions in absolute pitch ability. Proc. Natl. Acad. Sci. 104, 14795–14800 (2007).

30. Deutsch, D., Dooley, K., Henthorn, T. & Head, B. Absolute pitch among students in an American music conservatory: Association with tone language fluency. J. Acoust. Soc. Am. 125, 2398–2403 (2009).

31. Deutsch, D., Li, X. & Shen, J. Absolute pitch among students at the Shanghai Conservatory of Music: A large-scale direct-test study. J. Acoust. Soc. Am. 134, 3853–3859 (2013).

32. Bachem, A. Various Types of Absolute Pitch. J. Acoust. Soc. Am. 9, 146–151 (1937).

33. Van Hedger, S. C., Heald, S. L. M. & Nusbaum, H. C. The effects of acoustic variability on absolute pitch categorization: Evidence of contextual tuning. J. Acoust. Soc. Am. 138, 436–446 (2015).

34. Ross, D. A. & Marks, L. E. Absolute pitch in children prior to the beginning of musical training. Ann. N. Y. Acad. Sci. 1169, 199–204 (2009).

35. Ross, D. A., Olson, I. R., Marks, L. E. & Gore, J. C. A nonmusical paradigm for identifying absolute pitch possessors. J. Acoust. Soc. Am. 116, 1793–1799 (2004).

36. Ross, D. A., Gore, J. C. & Marks, L. E. Absolute pitch: Music and beyond. Epilepsy Behav. 7, 578–601 (2005).

37. Deutsch, D. 5 - Absolute Pitch. The Psychology of Music (Third Edition) (2013). doi:http://dx.doi.org/10.1016/B978-0-12-381460-9.00005-5

38. Dohn, A., Garza-Villarreal, E. A., Ribe, L. R. & Wallentin, M. Musical activity tunes up absolute pitch ability. Music Percept. 31, 359–371 (2014).

39. Bahr, N., Christensen, C. A. & Bahr, M. Diversity of accuracy profiles for absolute pitch recognition. Psychol. Music33, 58–93 (2005).

40. Miyazaki, K. & Ogawa, Y. Learning Absolute Pitch by Children. Music Percept. 24, 63–78 (2006).

